# The oncometabolite L-2-hydroxyglutarate is a common product of Dipteran larval development

**DOI:** 10.1101/2020.05.05.078857

**Authors:** Nader H. Mahmoudzadeh, Alexander J. Fitt, Daniel B. Schwab, William E. Martenis, Lauren M. Nease, Charity G. Owings, Garrett J. Brinkley, Hongde Li, Jonathan A. Karty, Sunil Sudarshan, Richard W. Hardy, Armin P. Moczek, Christine J. Picard, Jason M. Tennessen

## Abstract

The oncometabolite L-2-hydroxyglutarate (L-2HG) is considered an abnormal product of central carbon metabolism that is capable of disrupting chromatin architecture, mitochondrial metabolism, and cellular differentiation. Under most circumstances, mammalian tissues readily dispose of this compound, as aberrant L-2HG accumulation induces neurometabolic disorders and promotes renal cell carcinomas. Intriguingly, *Drosophila melanogaster* larvae were recently found to accumulate high L-2HG levels under normal growth conditions, raising the possibility that L-2HG plays a unique role in insect metabolism. Here we explore this hypothesis by analyzing L-2HG levels in 18 insect species. While L-2HG was present at low-to-moderate levels in most of these species (<100 pmol/mg; comparable to mouse liver), Dipteran larvae exhibited a tendency to accumulate high L-2HG concentrations (>100 pmol/mg), with the mosquito *Aedes aegypti,* the blow fly *Phormia regina,* and three representative *Drosophila* species harboring concentrations that exceed 1 nmol/mg – levels comparable to those measured in mutant mice that are unable to degrade L-2HG. Overall, our findings suggest that one of the largest groups of animals on earth commonly generate high concentrations of an oncometabolite during juvenile growth, hint at a role for L-2HG in the evolution of Dipteran development, and raise the possibility that L-2HG metabolism could be targeted to restrict the growth of key disease vectors and agricultural pests.

## INTRODUCTION

The field of cancer metabolism has become increasingly focused on how small molecule metabolites regulate cell proliferation and promote cancer progression (Martinez-Reyes and Chandel, 2020). In this regard, a number of compounds have emerged as oncometabolites – pro-growth molecules that enhance tumor growth by interfering with gene expression, epigenetic modifications, mitochondrial physiology, and signal transduction cascades (Mullen and DeBerardinis, 2012; Yang et al., 2013; Ye et al., 2018). These compounds, however, are not simply cancer-causing molecules, but also serve essential roles in normal metabolism and physiology. For example, the first molecules implicated as oncometabolites were the tricarboxylic acid intermediates fumarate and succinate (Raimundo et al., 2011), both of which play ancient and essential roles in energy production. Similarly, the oncometabolite D-2-hydroxyglutarate (D-2HG), which is perhaps best known for promoting glioblastoma (Ye et al., 2018), also serves normal metabolic roles in bacteria, yeast, and even humans (Becker-Kettern et al., 2016; Struys et al., 2005; Zhang et al., 2017). Such examples illustrate how oncometabolites act in diverse and important metabolic mechanism across all kingdoms of life and suggest that studying normal oncometabolite function can advance our understanding of how these molecules induce human disease.

Among known oncometabolites, the compound L-2HG stands out as unusual because eukaryotes lack enzymes dedicated to L-2HG production. Mammalian cells synthesize L-2HG as a result of promiscuous Lactate Dehydrogenase (Ldh) and Malate Dehydrogenase (Mdh) activity and degrade this compound via the enzyme L-2HG dehydrogenase (L2HGDH) (Ye et al., 2018). Most mammalian tissues, with the exception of the mouse testis, maintain low L-2HG concentrations and inappropriate L-2HG accumulation can induce dramatic changes in epigenetic modifications, central carbon metabolism, and growth factor signaling (Ma et al., 2017; Teng et al., 2016; Ye et al., 2018). As a result, nearly all studies of L-2HG focus on the detrimental effects of this compound in neurological disorders and renal cell carcinoma and the question remains as to whether L-2HG, like the other oncometabolites, serves a normal physiological role (Ma et al., 2017; Shim et al., 2014; Ye et al., 2018). In this regard, several studies have observed that cultured mammalian cells accumulate excess L-2HG in response to oxidative stress, with hypoxia, acidic pH, and elevated NADH levels enhancing L-2HG synthesis and accumulation (Intlekofer et al., 2015; Intlekofer et al., 2017; Mullen et al., 2014; Nadtochiy et al., 2016; Oldham et al., 2015; Reinecke et al., 2012; Teng et al., 2016). These cell culture studies suggest that L-2HG metabolism could act as a metabolic signaling molecule that helps cellular physiology adapt to redox stress; however, the functions of L-2HG function *in vivo,* if any, remain unknown.

One of the only examples of a healthy animal accumulating high L-2HG levels in a regulated and predictable manner is during larval development of the fruit fly *Drosophila melanogaster* (Li et al., 2017). While the exact reason for why *Drosophila melanogaster* larvae accumulate L-2HG remains to be elucidated, this observation raises the possibility that L-2HG serves a unique role in insects and suggests that comparative studies of insect metabolism could illuminate the endogenous function of this oncometabolite. Towards this goal, we used gas chromatography-mass spectrometry to quantify L-2HG levels in 18 species of insects. Our analysis revealed that representative species across the order Diptera seem particularly adept at accumulating very high L-2HG concentrations during larval development. Moreover, we demonstrate that while Dipteran larvae can generate excess L-2HG in response to hypoxia, larvae generate high concentrations of this compound regardless of oxygen concentration. This finding indicates that one of the largest and most diverse animal orders on the planet commonly produces high concentrations of an oncometabolite in a developmentally-regulated manner and suggests that further studies of Dipteran L-2HG metabolism could help elucidate the endogenous functions of this compound. In addition, our study suggests the L-2HG metabolism serves a unique role in the Dipteran development, thus raising the possibility that production of this compounds could be used to control populations of common disease vectors and agricultural pests.

## METHODS

### Insect Husbandry

#### Libellulida

(Odonata, Anisoptera, Libellulidae) species and *Enallagma* (Odonata, Zygoptera, Coenagrionidae) species.: Nymphs were purchased from Carolina Biological Supply *(Libellulida* species, 143526; *Enallagma* species, 143520) and maintained following the supplier’s recommendation. For collections, individual nymphs were removed from the culture, briefly patted dry, placed in a pre-tared 2 mL tube containing 1.4 mm ceramic beads, massed, and flash frozen in liquid nitrogen.

#### Gryllodes sigillatus

(Orthoptera, Gryllidae): Nymphs were obtained from Carolina Biology (item #143558). Upon arrival, cultures were maintained at ambient temperature and fed a diet of dried dog food, lettuce, and apple slices. All samples contained a single individual.

#### Oncopeltus fasciatus

(Hemiptera, Lygaeidae): Milkweed bugs were obtained from Carolina Biological Supply (Item # 143800) and maintained on organic sunflower seeds.

#### Apis mellifera

(Hymenoptera, Apidae): Bee samples were collected from a colony established and maintained by Dr. Irene Newton’s lab at Indiana University-Bloomington. Samples were collected in 2018 during the months of June, July, and August. For both larvae and adults, each sample contained an individual animal. Adult samples consisted of workers.

#### Onthophagus taurus

(Coleoptera, Scarabaeidae): Adults were collected from cow dung pads at Busselton, Western Australia (−33° 39’ 8” S, 115° 20’ 43” E) in January 2016 and shipped to Bloomington, IN, for rearing. All beetles were maintained as a single colony in the laboratory at 24°C on a 16 L: 8 D cycle, and fed homogenized cow dung *ad libitum* following an established protocol (Moczek et al., 2002). In order to obtain offspring, beetles were allowed to breed in plastic containers (25 cm tall × 20 cm in diameter) and filled ~75% with a moist sand:soil mixture. For each round of breeding, six female and three male beetles were added to one breeding container and provisioned with ~0.5 L of homogenized cow dung. Beetles were allowed to breed for one week, at which point they were recaptured and brood balls containing offspring were collected and placed in plastic containers. All samples contained a single individual.

#### Tribolium castaneum

(Coleoptera, Tenebrionidae): Cultures were maintained on King Arthur whole wheat flour supplemented with active dry yeast at 28°C and 55-65% humidty. For larval analysis, 3^rd^ and 4^th^ instar larvae were collected. For adult hypoxia experiment, recently emerged adults were collected and adults were kept on new flour for the duration of the treatment. The strain *Vermillion^white^* was used for all the studies in this report.

#### Tenebrio molitor

(Coleoptera, Tenebrionidae): Larvae (~100-200 mg; Item #144274 and ~20 mg; Item #144287) and adults (Item #144270) were purchased from Carolina Biological Supply and fed whole wheat flour and apple slices, as per the distributors care sheet. All samples contained an individual animal. For larvae of a mass >100 mg, individual animals were homogenized in 800 μl of methanol extraction buffer as described below (see *Sample collection and L-2HG Quantification*) and the homogenate was immediately diluted 1:2 or 1:4 with additional extraction buffer, depending on the mass of the larvae. All samples contained a single individual.

#### Galleria mellonella

(Lepidoptera, Pyralidae): Larvae were purchased from Carolina Biological Supply (Item # 143928) and raised on the provided media according to the distributors care sheet. All samples contained a single individual.

#### Vanessa cardui

(Lepidoptera, Nymphalidae): Larvae were obtained from Carolina Biology (Item # 144076) and maintained on the culture media provided by the distributor at ambient temperature. All samples contained a single individual.

#### Manduca sexta

(Lepidoptera, Sphingidae): Larvae were obtained from Carolina Biological Supply (#143886) and maintained at room temperature on hornworm diet (Carolina Biological Supply; #143910). Animals were collected throughout the L3 stage. Each sample consisted of an individual animal. For larvae with a mass greater than 50 mg, individual animals were homogenized in 800 μl of methanol extraction buffer as described below (see *Sample collection and L-2HG Quantification*) and the homogenate was immediately diluted 1:2 or 1:4 with additional extraction buffer, depending on the mass of the larvae.

#### Aedes aegypti

(Diptera, Nematocera, Culicidae): RexD (Puerto Rico) derived Higgs White Eye (HWE) strain were maintained in an insect incubator (Percival Model I-36VL, Perry, IA, USA) at 28°C and 75% relative humidity with a 12h:12h light:dark cycle. Larvae were reared in freshwater (dH2O) at a density of 200 larvae/L of water. Water was changed every other day. Each larval cup was fed a 4:1 mixture of finely ground fish pellets to baker’s yeast a day. Adult mosquitos were fed 10mL of 10% sucrose daily via cotton balls. Samples contained of 10-20 mg of third instar larvae.

#### Hermetia illucens

(Diptera, Brachicera, Stratiomyidae): Larval cultures were shipped from Dr. Jeffery Tomberlin’s lab (Texas A&M University; College Station, Texas, United States) and collected upon arrival. Each sample contained an individual larva. Cultures were maintained at ambient temperature and adults were collected upon emergence.

#### Musca domestica

(Diptera, Brachicera, Muscidae): Larvae were purchased from Carolina Biological Supply (Item #144410) and raised on Instant House Fly Medium (Item # 144424) at 25°C. Individual larva were collected in a 1.5 ml microfuge tube and immediately placed on ice. Samples were washed as described for the *Drosophila* species (see above). All samples contained a single individual.

#### Drosophila species

(Diptera, Brachycera, Drosophilidae): All *Drosophila* species were maintained on standard Bloomington *Drosophila* Stock Center (BDSC) media at 25°C. The *Drosophila melanogaster* strain *w^1118^* was used for all experiments. *Drosophila hydei* and *Drosophila busckii* cultures, which were kindly provided by Dr. Irene Newton’s lab (Indiana University-Bloomington, USA), are derived from wild isolates collected in Brown County, Indiana.

For all species, virgin males and females were collected immediately following eclosion and aged for 3 days on BDSC media prior to collection or treatment. For larval analyses, embryos were collected on molasses agar with covered with yeast paste as previously described (Li and Tennessen, 2018). Larvae were allowed to develop for 60 hours (*D. melanogaster* and *D. busckii)* or 84 hours (*D. hydei)* prior to collection. For hypoxic and hyperoxic treatments, larvae were placed in 35 mm plates with Whatman filter paper at the bottom that was wetted with 2 ml of phosphate buffer saline (PBS; pH 7.4) and contained approximately 500 mg of yeast paste in the center. Regardless of treatment or age, larval samples were collected by placing ~20 mg of larvae in a 1.5 ml microfuge tube on ice. Larvae were washed at least three times using ice-cold PBS to remove all yeast and debris from the sample. Following the final wash, all PBS was removed from the sample and the tube was flash frozen in liquid nitrogen.

#### Phormia regina

(Diptera, Brachicera, Calliphoridae): All flies were collected from a laboratory colony (> 5 generations) that was generated from wild-caught *P. regina* (collected from Military Park, Indianapolis, IN, USA) and maintained in a 30 × 30 × 30cm cage (Bioquip, Rancho Dominguez, CA) within the IUPUI “fly room.” The colony was reared at ~25°C ambient temperature, 60% ambient humidity and 24 hour light and were provided sugar and water ad libitum. Chicken liver was provided to the colony ~ 1 week post-eclosion for ovary maturation. 2-4 days following ovary maturation, chicken liver (25g) was provided as the egg oviposition substrate for a period of 4-6 hours. Following oviposition, the cup containing the chicken liver and eggs were placed in a one-quart glass jar half-filled with fine pine shavings (Lanjay Inc., Montreal, QC). Larvae were allowed to develop under ambient conditions. For sample collection, an individual third instar larva was placed in a 1.5 ml microfuge tube on ice. Samples were washed as described for the *Drosophila* species (see above). For adult analysis, adult flies were randomly collected 3 – 5 days post-emergence from the lab colony.

### Method for Mice Liver Harvest

Both male and female C57BL/6J mice aged 2-4 months were given normal chow (Labdiet) ad libitum. For mice starvation, chow was removed in the evening 14 hours prior to tissue harvest. At time of harvest, mice were anesthetized using isoflourane (Vetone) and blood was collected. Mice were then euthanized. Tissues were briefly washed in chilled DPBS (Corning), dried using kimwipes (Kimberly-Clark) and snap frozen using liquid nitrogen. All animal studies were approved by institutional animal care and use committee (IACUC).

### Sample collection and L-2HG Quantification

For pooled samples that contained multiple individuals, samples were collected in 1.5 ml microfuge tubes and flash frozen in liquid nitrogen. Prior to homogenization, tubes were removed from liquid nitrogen, the cap end swiftly pounded against the desktop to dislodge the sample, and the pellet was transferred into a pre-tared 2 ml screwcap tube containing 1.4 mm ceramic beads. The sample mass was immediately measured with an analytical balance, the tube was flash frozen in liquid nitrogen, and samples were stored at −80°C. For samples that contained an individual insect, the animal was collected and immediately transferred into a pre-tared 2 ml screwcap tube containing 1.4 mm ceramic beads. The sample mass was measured using an analytical balance, the tube was flash frozen in liquid nitrogen, and samples were stored at −80 °C.

Homogenization and metabolite extractions were conducted as previously described (Li and Tennessen, 2019). Briefly, samples were removed from the −80°C freezer and placed into a benchtop enzyme cooler pre-chilled to −20°C. 800 μl of prechilled methanol extraction buffer (−20°C; 90% methanol diluted with HPLC-grade water + 8 μg (RS)-2-Hydroxy-1,5-pentanedioate-2,3,3-d3) was added to individual sample tubes. Sample were homogenized in an Omni Beadruptor 24. Homogenized samples were returned to a −20°C benchtop enzyme cooler and incubated in a −20°C freezer for one hour. Samples were then centrifuged at ~20,000 × g in a refrigerated centrifuge for 5 minutes to pellet insoluble debris. 600 μl of the supernatant was removed and transferred to a 1.5 ml microfuge tube and dried overnight in a vacuum centrifuge. Samples were then derivatized using a two-step method involving R-2-butanol and acetic anhydride.

Derivatized samples were injected (1.5 μL, 1:5 split ratio) into an Agilent GC6890-5973i instrument using a Gerstel MPS autosampler. Separation of compounds was achieved by gas chromatography (GC) with a Phenomex ZB5-5 MSi column. The GC was programmed to increase temperature as follows: (1) Inlet temperature was set to 250 °C and initial temperature was set to 95 °C with a one-minute hold. (2) Temperature was increased at a rate of 40 °C per minute until it reached 110 °C with a two-minute hold. (3) Temperature was increased a rate of 5 °C per minute to 250 °C. (4) Temperature was increased at a rate of 25 °C per minute to 330 °C followed by a three-minute hold. Selected ion monitoring (SIM mode) was programmed to record m/z ion values 173 for endogenous D-/L-2HG, and 176 for the deuterated D-/L-2HG internal heavy standard. Concentration in each sample was calculated by comparison to the internal standard and normalized to the sample mass.

### Hypoxia and Hyperoxia Treatments

For all manipulations of atmospheric oxygen concentration, samples were placed in an air sealed plexiglass chamber equipped with a pressure release valve that was located within a 25°C temperature-controlled room. A Sable systems ROXY-4 gas regulator was used to control oxygen concentration within the chamber. Desired oxygen concentrations were maintained by the ROXY-4 system by injecting either N_2_ or O_2_ gas into the chamber.

### Statistical Analysis

All data analyses were conducted using JMP v. 14. Prior to analysis, variables were evaluated for normality and homoscedasticity using Shapiro-Wilk and Levene’s tests, respectively. Where these assumptions were met, we used two-tailed pooled t-tests or ANOVA followed by Tukey-Kramer HSD test to determine differences among treatment groups. Where these assumptions were not met, we conducted non-parametric Wilcoxon rank sum or Kruskal-Wallis tests. To compare L-2HG titers across larval insect species with mouse liver samples (see Fig. 1A), we twice conducted Dunnett’s post hoc test with a Bonferroni adjustment for multiple comparisons.

**Figure 1.**
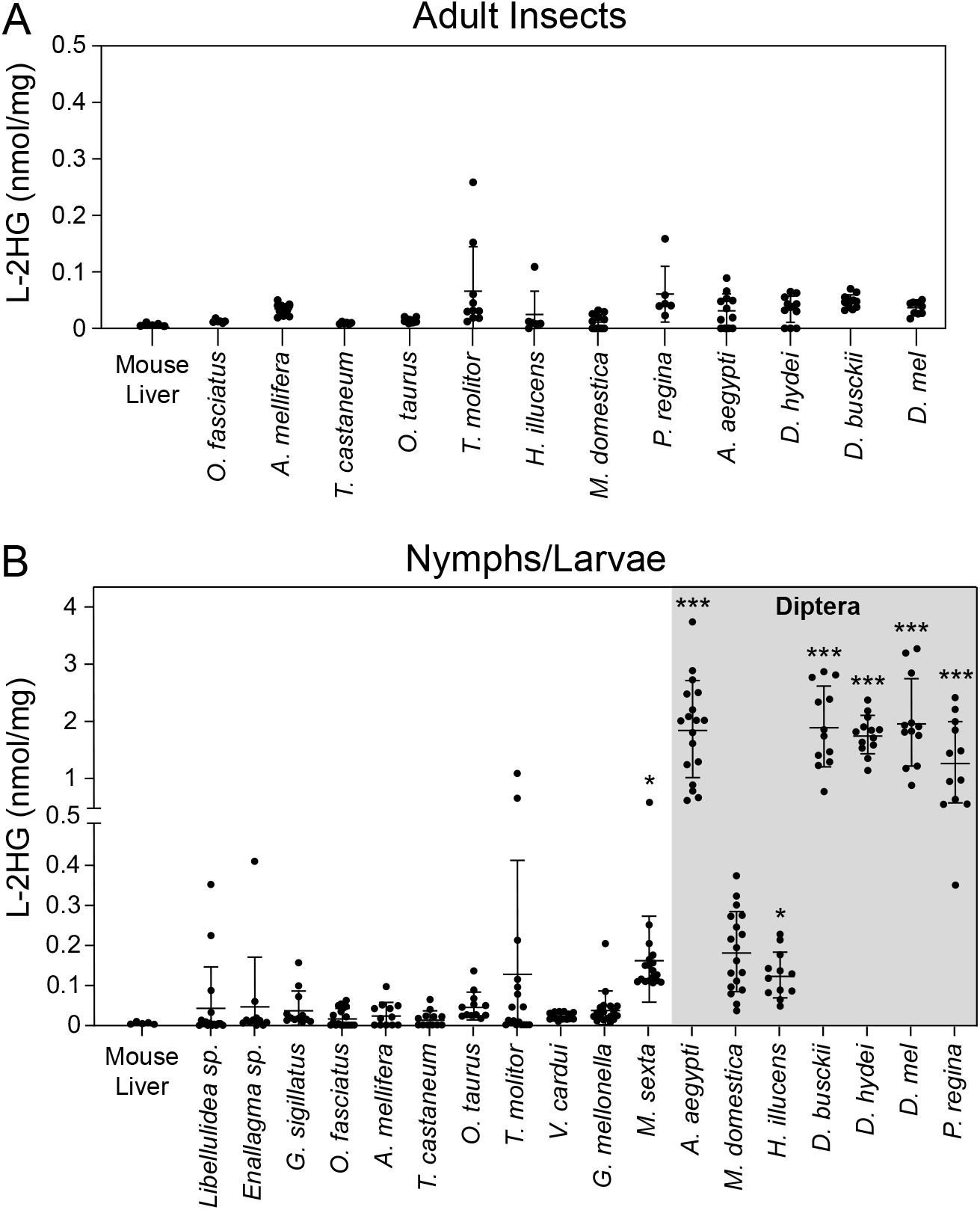
Dipteran larvae accumulate high L-2HG levels. L-2HG levels were measured in (A) adult insects, (B) juvenile insects, and (A,B) mouse liver, which served as a baseline control. Asterisks indicate that L-2HG levels are significantly higher than those measured in mouse liver. Data are presented in scatter plots with mean ± SD. *P<0.05; ***P<0.001. See Supplemental Methods for a description of the statistical analysis.

## RESULTS AND DISCUSSION

Recent findings that *D. melanogaster* larvae accumulate high L-2HG levels motivated us to determine if this molecule is abundant in other insects. Towards this goal, we used a chiral derivatization method coupled with gas chromatography-mass spectrometry (GC-MS) to quantify L-2HG levels in a diversity of insect species and mouse liver, which is known to possess low L-2HG levels and served as a baseline control (Ma et al., 2017). Among those animals and developmental stages surveyed, we observed no significant difference in L-2HG levels between mouse liver and any adult insect (Figure 1A). Similarly, L-2HG levels were present at a low level in nymphal stages of four hemimetabolous species (i.e., insects that do not undergo complete metamorphosis; Figure 1B) and larval stages of holometabolous species including the European honey bee *(Apis mellifera),* three species of Coleoptera (beetles, *Tribolium castanaeum, Tenebrio molitor,* and *Onthophagus taurus),* and two Lepidopteran (moths and butterflies) species *(Vanessa cardui* and *Galleria mellonella).* Meanwhile, third instar larvae of the moth *Manduca sexta* accumulated L-2HG levels that were slightly, but significantly, higher than mouse liver (Figure 1B) – a result that was observed throughout the entire third instar, independent of body mass (Figure S1).

In contrast to these insects, a majority of the Dipteran species examined harbored notably elevated larval L-2HG levels, with the mosquito *Aedes aegypti,* the blow fly *Phormia regina,* and multiple members of genus *Drosophila (D. melanogaster, D. busckii,* and *D. hydei)* accumulating L-2HG levels that exceeded 1 nmol/mg. These L-2HG concentrations are comparable to those measured in both humans and mice lacking the enzyme L-2-hydroxyglutarate dehydrogenase, which is responsible for degrading L-2HG (Ma et al., 2017; Rzem et al., 2004). In fact, of the seven Dipteran species examined in this study, only the house fly, *Musca domestica,* maintained larval L-2HG levels that were not significantly higher than those observed in mouse liver tissue (Figure 1B); however, we note that even these *M. domestica* samples contained an average of >100 pmol/mg. Overall, our observations suggest that Dipterans exhibit a unique tendency to accumulate L-2HG during larval development.

Dipteran larvae develop in moist environments. Since human cells accumulate L-2HG in response to hypoxia and decreased electron transport chain activity (Intlekofer et al., 2015; Mullen et al., 2014; Oldham et al., 2015), we examined the possibility that the Dipteran species analyzed herein accumulate high L-2HG levels as the result of growing within a potentially hypoxic environment (e.g., yeast paste, water, and rotting chicken liver). To test this hypothesis, we first determined if hypoxia is capable of inducing L-2HG accumulation in adult males of these species, which normally harbor low L-2HG levels (Figure 1A). Consistent with studies of mammalian cells, adult male *Aedes aegypti, Drosophila melanogaster, Drosophila hydeii,* and *Phormia regina* accumulated excess L-2HG when exposed to 1% O_2_ (~1 kPa O_2_) for 6 hours (Figure 2A). We observed a similar phenomenon in third instar larvae of these same species where a 6 hour exposure to 1% O_2_ (an O_2_ level that is incompatible with *Drosophila melanogaster* larval development, Zhou et al., 2008) generated significantly higher L-2HG levels than normoxic controls (~20 kPa O_2_; Figure 2B). In fact, several individual blowfly larvae harbored L-2HG concentrations that exceeded 6 nmol/mg, which are among the highest levels ever recorded in animal tissues, even exceeding the L-2HG levels observed in *L2hgdh* mutant mouse brains and testis (Ma et al., 2017). These observations demonstrate that, similar to mammalian cells, Dipterans appear capable of accumulating L-2HG in response to hypoxia and are consistent with recent observations that *Drosophila melanogaster* generates L-2HG in response to artificially induce mitochondrial stress (Hunt et al., 2019).

**Figure 2.**
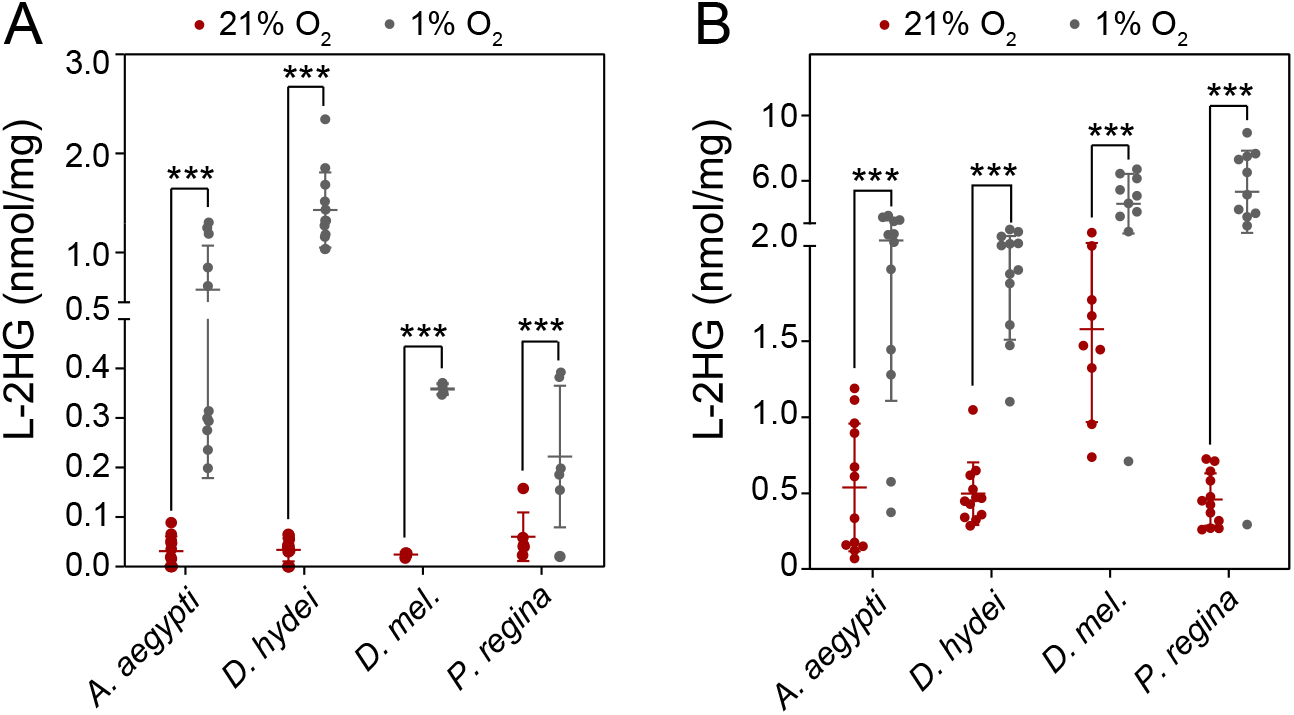
Dipteran insects accumulate L-2HG in response to hypoxia. L-2HG levels were measured in select Dipteran (A) adults or (B) larvae following a 6-hour incubation in the presence of 1% O_2_. Data are presented in scatter plots with mean ± SD. ***P<0.001. See Supplemental Methods for a description of the statistical analysis.

Our findings that a low oxygen environment results in elevated L-2HG levels raises the question as to whether the high L-2HG concentrations observed in Dipteran larvae is simply the result of transient exposure to hypoxia. We tested this possibility by measuring L-2HG levels in larvae following a 24-hour exposure to mild hypoxia (5%, an O_2_ level that delays, but does not arrest larval development, Zhou et al., 2008), normoxia, or hyperoxia (30% O_2_, 50% O_2_). If the high L-2HG levels observed in these larvae simply result of hypoxic stress, we would expect that L-2HG levels would inversely correlated with O_2_ concentration. Instead, we observe that species-specific L-2HG concentrations remained largely unchanged following 24-hour exposure to any of these four oxygen concentrations (Figure 3), suggesting that oxygen availability is not the primary driving force behind L-2HG accumulation in these animals. Moreover, our finding that 1 % O_2_, but not 5 % O_2_, induces excess L-2HG accumulation supports previous observations in *Drosophila melanogaster* that larvae mount different physiological responses depending on the severity of hypoxic conditions (Lavista-Llanos et al., 2002).

**Figure 3.**
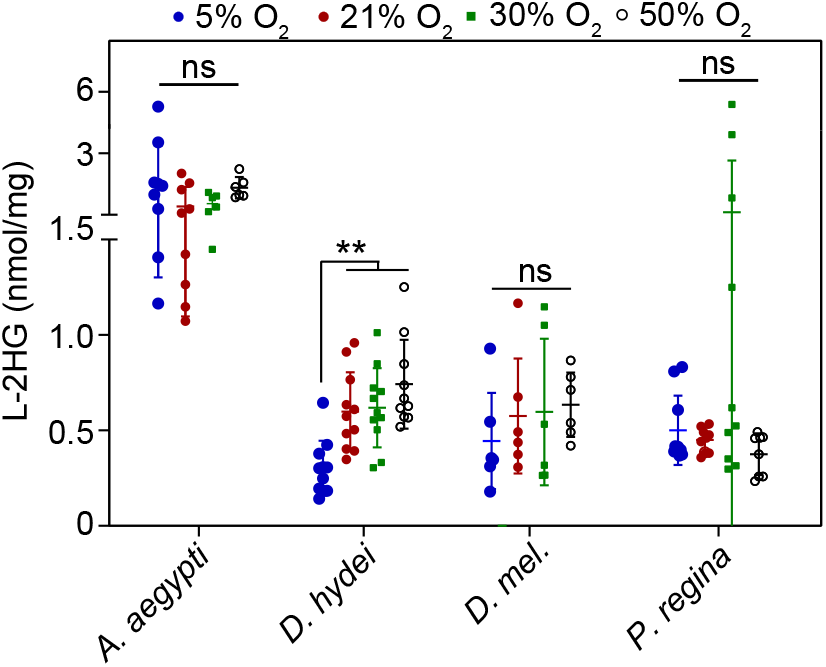
Larval L-2HG levels do not change in response to mild hypoxia or hyperoxia. L-2HG levels were measured in select Dipteran larvae following a 24-hour incubation in the presence of 5%, 21%, 30% or 50% O_2_. Data are presented in scatter plots with mean ± SD. **P<0.01; See Supplemental Methods for a description of the statistical analysis.

Overall, our results demonstrate that several Dipteran species accumulate high levels of the oncometabolite L-2HG during normal larval development. While the endogenous functions of L-2HG within these insects remains to be elucidated, our observations raise important considerations. Insects are among the most diverse groups of animals on the planet, display complex life histories, and are adaptable to a wide-range of environmental conditions. When considered in this context, our survey of L-2HG metabolism is small in both the number of species and life-stages surveyed. Despite this limitation, we uncovered several instances where larvae generated relatively high L-2HG concentrations – a result which implies that this compound is not simply an oncometabolite or a waste product of the TCA cycle, but rather accumulates during normal development of a potentially large number of animal species.

The amount of L-2HG found within Dipteran larvae is striking, as similar L-2HG concentrations in mammals are associated with severe neurometabolic defects and renal tumors (Shim et al., 2014; Ye et al., 2018), suggesting that this compound serves a unique role in Dipteran physiology when compared to other animals. One explanation for our observations is that developmentally regulated L-2HG accumulation acts as part of a metabolic program that protects Dipteran larvae from transient exposure to extreme hypoxia. L-2HG has been repeatedly observed to be produced in animal cells exposed to hypoxic conditions and the production of this molecule is thought to play a role in the cellular hypoxia response. Moreover, Dipteran larvae have evolved to be exceptionally tolerant of hypoxia and anoxia, as evident by the ability of *D. melanogaster* larvae to remain motile for over 30 minutes under anoxic conditions (Callier et al., 2015). Based on these observations, we propose that L-2HG accumulation helps Dipteran development survive transient exposure to hypoxic conditions that may be common in the larval environment.

Our findings also raise the question as to how the Dipteran species analyzed in this study have evolved to tolerate such high L-2HG levels. L-2HG is a potent inhibitor of enzymes that use α-ketoglutarate (α-KG) as a substrate and high concentrations of this molecule interfere with a diversity of α-KG-dependent processes, which include mitochondrial metabolism, the removal of methyl groups for DNA and histones, and stabilization of the transcription factor HIF1α. Considering that the L-2HG concentration observed in *Aedes aegypti*, *Phormia regina*, and the three *Drosophila* species used in this study exceeds all previous reported Ki values for α-KG-dependent enzymes, the cellular physiology of these systems must be uniquely adapted to the presence of this oncometabolite. Future studies should examine how Dipterans have evolved to tolerate concentrations of this compound that would prove fatal to humans.

The precise endogenous L-2HG functions notwithstanding, our study highlights how the natural diversity of insects remains a remarkable resource for discovering and exploring the metabolic mechanisms that support animal growth. In addition, these findings demonstrate how studying Dipteran development can identify unique metabolic features that could be targeted for controlling both agricultural pests and human disease vectors.

## ACKNOWLEDGEMENTS

We thank the Newton, Zelhof, and Tomberlin labs for providing for samples and advice. S.S. is supported by NCI R01CA200653. G.B. is supported by NCI F30CA232397. A.P.M. is supported by NSF 1901680. J.A.K. and JMT are supported by NSF 1726633. J.M.T. is supported by NIH 1R35GM119557.

## Supplemental Figure Legend

**Figure S1.**
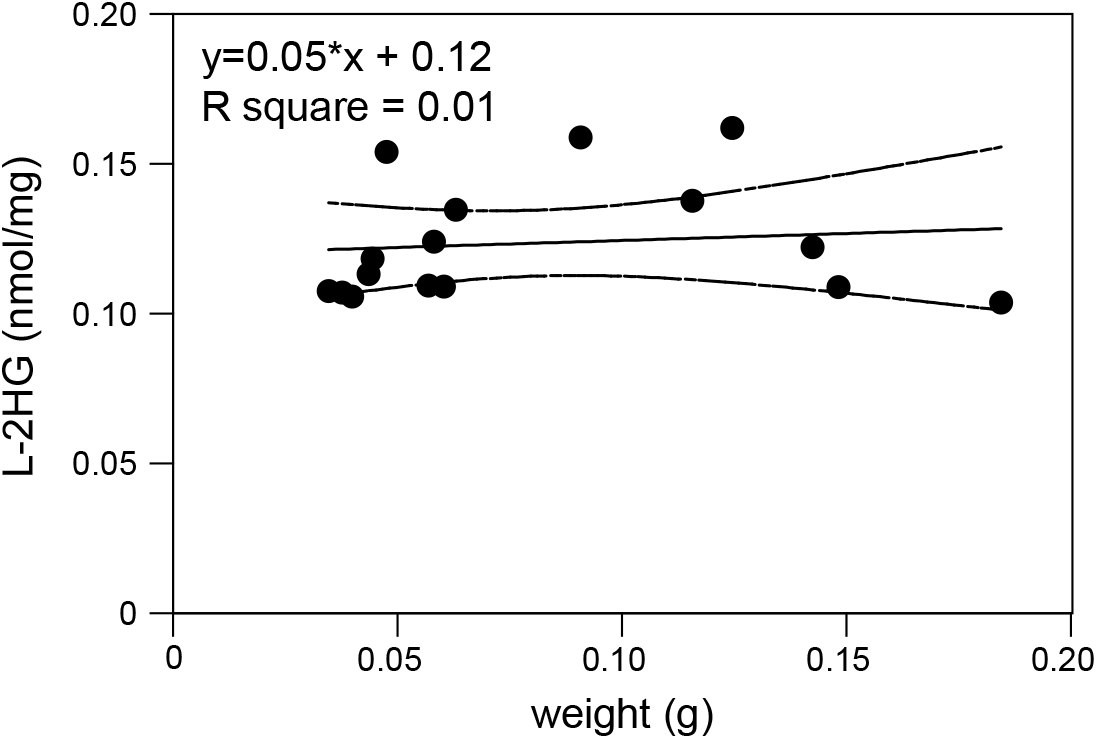
Concentration of L-2HG relative to body mass in *M. sexta* third instar larvae. Data are presented as a scatter plot. Each data point represents an individual larva. Dashed lines represent the 95% confidence interval.

## LITERATURE CITED

Becker-Kettern, J., Paczia, N., Conrotte, J. F., Kay, D. P., Guignard, C., Jung, P. P. and Linster, C. L. (2016). *Saccharomyces cerevisiae* Forms D-2-Hydroxyglutarate and Couples Its Degradation to D-Lactate Formation via a Cytosolic Transhydrogenase. J Biol Chem 291, 6036–6058.

Callier, V., Hand, S. C., Campbell, J. B., Biddulph, T. and Harrison, J. F. (2015). Developmental changes in hypoxic exposure and responses to anoxia in Drosophila melanogaster. J Exp Biol 218, 2927–2934.

Hunt, R. J., Granat, L., McElroy, G. S., Ranganathan, R., Chandel, N. S. and Bateman, J. M. (2019). Mitochondrial stress causes neuronal dysfunction via an ATF4-dependent increase in L-2-hydroxyglutarate. J Cell Biol 218, 4007–4016.

Intlekofer, A. M., Dematteo, R. G., Venneti, S., Finley, L. W., Lu, C., Judkins, A. R., Rustenburg, A. S., Grinaway, P. B., Chodera, J. D., Cross, J. R., et al. (2015). Hypoxia Induces Production of L-2-Hydroxyglutarate. Cell Metabolism.

Intlekofer, A. M., Wang, B., Liu, H., Shah, H., Carmona-Fontaine, C., Rustenburg, A. S., Salah, S., Gunner, M. R., Chodera, J. D., Cross, J. R., et al. (2017). L-2-Hydroxyglutarate production arises from noncanonical enzyme function at acidic pH. Nature chemical biology 13, 494–500.

Lavista-Llanos, S., Centanin, L., Irisarri, M., Russo, D. M., Gleadle, J. M., Bocca, S. N., Muzzopappa, M., Ratcliffe, P. J. and Wappner, P. (2002). Control of the hypoxic response in Drosophila melanogaster by the basic helix-loop-helix PAS protein similar. Mol Cell Biol 22, 6842–6853.

Li, H., Chawla, G., Hurlburt, A. J., Sterrett, M. C., Zaslaver, O., Cox, J., Karty, J. A., Rosebrock, A. P., Caudy, A. A. and Tennessen, J. M. (2017). *Drosophila* larvae synthesize the putative oncometabolite L-2-hydroxyglutarate during normal developmental growth. Proc Natl Acad Sci U S A 114, 1353–1358.

Li, H. and Tennessen, J. M. (2018). Preparation of Drosophila Larval Samples for Gas Chromatography-Mass Spectrometry (GC-MS)-based Metabolomics. J Vis Exp.

Li, H. and Tennessen, J. M. (2019). Quantification of D-and L-2-Hydroxyglutarate in Drosophila melanogaster Tissue Samples Using Gas Chromatography-Mass Spectrometry. Methods Mol Biol 1978, 155–165.

Ma, S., Sun, R., Jiang, B., Gao, J., Deng, W., Liu, P., He, R., Cui, J., Ji, M., Yi, W., et al. (2017). L2hgdh Deficiency Accumulates l-2-Hydroxyglutarate with Progressive Leukoencephalopathy and Neurodegeneration. Mol Cell Biol 37.

Martinez-Reyes, I. and Chandel, N. S. (2020). Mitochondrial TCA cycle metabolites control physiology and disease. Nat Commun 11, 102.

Moczek, A. P., Hunt, J., Emlen, D. J. and Simmons, L. W. (2002). Threshold evolution in exotic populations of a polyphenic beetle. Evol Ecol Res 4, 587–601.

Mullen, A. R. and DeBerardinis, R. J. (2012). Genetically-defined metabolic reprogramming in cancer. Trends Endocrinol Metab 23, 552–559.

Mullen, A. R., Hu, Z., Shi, X., Jiang, L., Boroughs, L. K., Kovacs, Z., Boriack, R., Rakheja, D., Sullivan, L. B., Linehan, W. M., et al. (2014). Oxidation of alpha-ketoglutarate is required for reductive carboxylation in cancer cells with mitochondrial defects. Cell Rep 7, 1679–1690.

Nadtochiy, S. M., Schafer, X., Fu, D., Nehrke, K., Munger, J. and Brookes, P. S. (2016). Acidic pH Is a Metabolic Switch for 2-Hydroxyglutarate Generation and Signaling. J Biol Chem 291, 20188–20197.

Oldham, W. M., Clish, C. B., Yang, Y. and Loscalzo, J. (2015). Hypoxia-Mediated Increases in l-2-hydroxyglutarate Coordinate the Metabolic Response to Reductive Stress. Cell Metab.

Raimundo, N., Baysal, B. E. and Shadel, G. S. (2011). Revisiting the TCA cycle: signaling to tumor formation. Trends Mol Med 17, 641–649.

Reinecke, C. J., Koekemoer, G., van der Westhuizen, F. H., Louw, R., Lindequie, J. Z., Mienie, L. J. and Smuts, I. (2012). Metabolomics of urinary organic acids in respiratory chain deficiencies in children. Metabolomics 8, 264–283.

Rzem, R., Veiga-da-Cunha, M., Noel, G., Goffette, S., Nassogne, M. C., Tabarki, B., Scholler, C., Marquardt, T., Vikkula, M. and Van Schaftingen, E. (2004). A gene encoding a putative FAD-dependent L-2-hydroxyglutarate dehydrogenase is mutated in L-2-hydroxyglutaric aciduria. Proc Natl Acad Sci U S A 101, 16849–16854.

Shim, E. H., Livi, C. B., Rakheja, D., Tan, J., Benson, D., Parekh, V., Kho, E. Y., Ghosh, A. P., Kirkman, R., Velu, S., et al. (2014). L-2-Hydroxyglutarate: an epigenetic modifier and putative oncometabolite in renal cancer. Cancer Discov 4, 1290–1298.

Struys, E. A., Verhoeven, N. M., Ten Brink, H. J., Wickenhagen, W. V., Gibson, K. M. and Jakobs, C. (2005). Kinetic characterization of human hydroxyacid-oxoacid transhydrogenase: relevance to D-2-hydroxyglutaric and gamma-hydroxybutyric acidurias. J Inherit Metab Dis 28, 921–930.

Teng, X., Emmett, M. J., Lazar, M. A., Goldberg, E. and Rabinowitz, J. D. (2016). Lactate Dehydrogenase C Produces S-2-Hydroxyglutarate in Mouse Testis. ACS Chem Biol 11, 2420–2427.

Yang, M., Soga, T. and Pollard, P. J. (2013). Oncometabolites: linking altered metabolism with cancer. J Clin Invest 123, 3652–3658.

Ye, D., Guan, K. L. and Xiong, Y. (2018). Metabolism, Activity, and Targeting of D-and L-2-Hydroxyglutarates. Trends Cancer 4, 151–165.

Zhang, W., Zhang, M., Gao, C., Zhang, Y., Ge, Y., Guo, S., Guo, X., Zhou, Z., Liu, Q., Zhang, Y., et al. (2017). Coupling between d-3-phosphoglycerate dehydrogenase and d-2-hydroxyglutarate dehydrogenase drives bacterial l-serine synthesis. Proc Natl Acad Sci U S A 114, E7574–E7582.

Zhou, D., Xue, J., Lai, J. C., Schork, N. J., White, K. P. and Haddad, G. G. (2008). Mechanisms underlying hypoxia tolerance in Drosophila melanogaster: hairy as a metabolic switch. PLoS Genet 4, e1000221.

